# A novel mobile genetic element with virus-like characteristics is widespread in the world’s oceans

**DOI:** 10.1101/2025.08.03.668216

**Authors:** Paul C. Kirchberger, Antoni Luque

## Abstract

While the majority of viruses falls within a handful of well-defined realms, many novel viruses and virus-like genetic elements remain to be discovered. Given the abundance and diversity of small, circular DNA genomes in animal gut viromes, we hypothesized that viruses of similar size might have been overlooked in other environments. Re-analyzing existing ocean virome datasets for small viruses, we discover a novel group of circular mobile genetic elements of ∼4.6kb in size, likely with double-stranded DNA, which we term Charybdis elements. Charybdis elements form a singular group based on cohesive gene content, encode around 10 hypothetical genes but apparently contain no virus, transposon or plasmid associated hallmark proteins. They are detected exclusively in marine datasets ranging from surface water to hydrothermal vents and sediments, from the Arctic to the Antarctic. Virome datasets separating virus-like particles by density gradients show that Charybdis elements accumulate in the fraction representing extracellular vesicles and small, tail-less viruses that are often associated with lipid membranes. One specific clade of Charybdis elements is shown to integrate into the genomes of the common marine group II archaeon candidate order Poseidoniales, the putative host of these elements. Analysis of thousands genomes shows a common genome architecture consisting of two modules: An accessory module containing a diverse range methyltransferases and other genes likely involved in host interactions, and a more conserved (putative) structural module. The structural module encodes proteins that are confidently predicted to form striking tri- or pentameric structures. These structures show stark resemblance and in some cases homology to protrusion or tail-elements on the capsids of viruses from the realms *Varidnaviria* and *Duplodnaviria*. Given these traits, Charybdis elements likely represent a hitherto undiscovered group of virus or virus-satellites, with at least one protein of a novel double-Greek-key fold serving as a potential capsid element. However, since their hosts remain unculturable, the true nature of these ubiquitous marine elements remains as of yet unresolvable.

## Introduction

Fundamentally, a virus is a mobile genetic element (MGE) composed of two parts: (I) a structural module encoding one or more proteins involved in building a protective virus shell (the capsid), and (II) a replication module encoding proteins such as DNA or RNA polymerases involved in copying the viral genome (1). Based on the presence of specific hallmark proteins, the virosphere is currently separated into seven realms - examples are the double-jelly-roll fold capsids of the realm *Varidnaviria* or the palm-fold RNA dependent RNA polymerases/reverse transcriptases for the realm *Riboviria*. The seven realms cover the vast majority of the known virosphere and are of presumably independent evolutionary origins (1, 2). In addition to bona-fide viruses, a veritable zoo of MGEs with various degrees of resemblance to viruses, often but not always containing clear structural or replication modules [for example (3–6)], can be considered parts of the extended virosphere (7).

Viruses infecting archaea have the fewest representatives among most realms (8). They consist predominantly of I) ubiquitous viruses with double-stranded DNA genomes whose icosahedral capsids are adorned with elaborate tails that facilitate attachment and infection of hosts (realm *Duplodnaviria*) (9) II) comparatively rare non-tailed viruses of the diverse realms *Varidnaviria* and *Singelavaria* (10), III) the archaeal-specific filamentous *Adnaviria* (11), and IV) the filamentous single-stranded DNA viruses of the *Pleolipoviridae* within the realm *Monodnaviria* (12). However, many other archaeal viruses with either single or double-stranded DNA genomes appear distantly or unrelated to viruses from these realms (13–22). This number of oddball viruses within the archaea, coupled with equally unusual MGEs [e.g.(23–26)] hints at the presence of additional novel virus and virus-like elements in this domain of life.

In the age of metagenomics, most viruses are not physically isolated anymore, but rather they are detected through remote homology searches for hallmark proteins in sequencing datasets [for example (27–29)]. Viruses encoding genes that have mutated beyond the reach of the most sensitive detection methods, or those lacking hallmark genes in the first place, will often remain part of the unrecognizable viral “dark matter” (30). As an alternative to developing ever more sensitive detection methods to find more and more remote viral homologs, other strategies to discover novel viruses and MGEs need to be applied. For example, one could devise strategies to find virus genomes that are possible in theory, but have yet to be found. Examples of these are viruses linking realms, such as cruciviruses with elements of Monodnaviria and Riboviria (31) or mirusviruses with elements of Varidnaviria and Duplodnaviria (32). Additionally, one could target conspicuous gaps in genome size or possible capsid architectures, as hypothesized in miniature tailed phages (33)).

Based on the clear size dichotomy between large, tailed phages with dsDNA genomes and small ssDNA microviruses in the human gut (34), combined with the relative absence of the latter in marine environments (35), we hypothesized that additional small viruses of around 5kb (such as the *Obscuriviridae* of the *Varidnaviria* (36, 37)) should exist in the world’s oceans. Re-analyzing small, circular metagenomically assembled genomes, we discover a cohesive group of MGEs that is globally distributed in marine environments, contains what appears to be virus-tail-like structural genes but neither a recognizable gene involved in replication, nor a known realm-defining hallmark protein. Patterns of presence and absence in metagenomic datasets produced through different sequencing/sampling methodology indicate that these elements have double-stranded DNA genomes, appear enclosed in either small capsids or extracellular vesicles, and replicate via rolling-circle replication. Structural prediction of the proteins encoded in these small genomes hint at the presence of either tri- or pentameric protrusions resembling those of tailed *Duplodnaviria* or non-tailed *Varidnaviria*, respectively. We also identify a putative hallmark protein composed of a double beta-sandwich fold unrelated to known viral hallmarks. The elements form several divergent phylogenetic clades, one of which contains members that are integrated into the genomes of the (as of yet) unculturable Candidate Marine Group II archaeal order Poseidoniales (38). Based on their exclusively marine distribution and the whirlpool-like appearance of their Greek-key hallmark proteins, we tentatively name these widespread virus-like MGEs Charybdis elements, after the mythical Greek sea creature & daughter of Poseidon. At this stage, it is not possible to ascertain the true nature of these mobile genetic elements, but the combination of traits exhibited by them marks them as a novel member of the virosphere, warranting further exploration.

## Results

### Metagenomic discovery of a marine mobile genetic element encoding a novel double-greek key fold protein

In an effort to detect novel viruses with small genomes, we re-analyzed small (<15kb) circular MAGs from an arbitrarily chosen Mediterranean Sea virome dataset [PRJEB30684,(39)] that had been obtained via multiple displacement amplification, a metagenomic sample preparation method that enriches for small, circular DNA elements (40). A genome of 4.54 kb in size, with two groups of oppositely oriented open reading frames (which we termed gp1 to gp10), piqued our interest (**Fig. 1A**). Apart from a methyltransferase gene ubiquitously found in prokaryotes and their MGEs (41), only hits to hypothetical genes of ∼4.5kb metagenomic contigs, spuriously binned to a range of archaea and bacteria as well as similar genomes annotated as “circular genetic element” (deposited by Tisza et al. 2020 (28)) could be produced from amino acid homology searches (**Table S1)**. Prediction of protein structures yielded two hypothetical proteins of interest (**Fig. 1A, Fig. S1A**): Gp5, predicted as a single jelly-roll fold protein (SJR, ∼100aa in length) and Gp3 (∼230aa in length), which is predicted to assume a two-domain structure: Each domain is composed of a β-sandwich of seven β-sheets, including a characteristic Greek-key motif, connected by a loop region. Through iterative homology searches for Gp3, we generated a hidden Markov model which ultimately detected 710 Gp3-positive contigs with a median size of 4.3 kb in the viral but not cellular fractions from the same metagenomic study, in depth profiles from 15 to 2000m (**Fig. S1B**).

**Figure 1:**
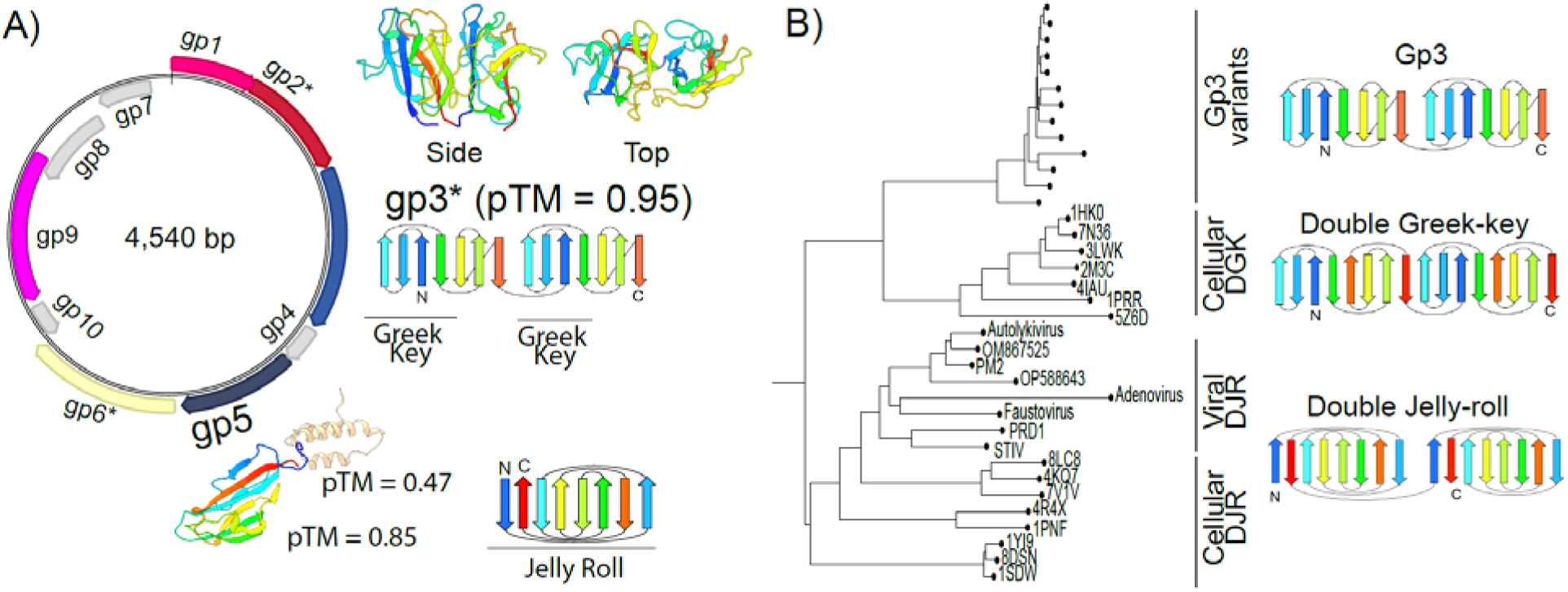
A circular MAG encoding a double Greek-key-like and a single jelly-roll protein. A) Exemplary genomic map and structural predictions for genes of interest. Arrows indicate genes. Rainbow colors on predicted protein structures correspond to beta-sheets depicted in the fold diagrams. Lines under fold diagrams indicate specific motifs. Alphafold3 pTM score represents the predicted template modeling confidence, with > 0.5 suggesting that the predicted complex might be similar to the true structure B) Structural phylogeny of exemplary Gp3 variants (see Table 1) with cellular/viral double Greek-key (DGK) and double jelly-roll fold (DJR) proteins. Neighbor-joining using the local distance difference test metric (LDDT) implemented in Foldtree (45). PDB IDs for experimentally resolved structures of cellular proteins are indicated in tree. Names of viruses with DJR capsids are written out and use available PDB IDs. PRD1 = 1cjd, PM2 = 2vvf, STIV = 2bbd, *Adenovirus* = 3tg7, *Faustovirus* = 5j7o. Other structures were predicted using Alphafold3 based on Gp3 sequences and major capsid proteins of NC_055066 (*Vibrio* phage 1.080.O._10N.286.48.A4, Autolykivirus), OP588643 (Phage Jorvik), OM867525 (vB_VruC_PG21).

No meaningful results could be attained from structural searches for homologs of Gp3 using Foldseek (42), with high confidence/e-value hits to a variety of jelly-roll domain proteins from all domains of life, despite the clear topological differences between jelly-rolls and the significantly older of Greek key motifs (43) (see Figure 1A). More sensitive Dali (44) searches resulted in medium-confidence hits for both double jelly-roll (e.g. 1PNF, a cellular amidase, Z-score: 5.6) and double Greek-key motif (e.g. 4IAU, a cellular beta-gamma crystallin, Z-score: 4.9) proteins. As Gp3 superficially resembles the double jelly-roll (DJR) fold hallmark proteins of the *Varidnaviria*, whose capsids also contain SJR proteins, we initially hypothesized that genomes carrying Gp3 might be highly divergent members of this viral realm. To investigate the homology between their hallmark proteins, we constructed a structural phylogeny of Gp3, double Greek-key and DJR proteins. However, the Gp3 proteins do not fall into any known structural cluster, indicating that they are neither capsid proteins of the *Varidnaviria* nor classic cellular Greek-key proteins (**Fig.1B**). Due to the marine nature of these MGEs and the whirlpool-like appearance of their Greek-key motifs when viewed from the top, we tentatively name these genomes Charybdis-elements.

### Global distribution and detection in low-density fractions of viromes

To investigate the broader distribution of Charybdis-elements, we queried the Ocean Gene Atlas web interface (46) for Gp3 proteins. Gp3 showed ubiquitous presence in the TARA Oceans Microbiome Reference Gene Catalog vI and vII datasets (47, 48) (Fig. 2A). As with the initial Mediterranean Sea dataset (**Fig. S1B**), detection succeeded in a variety of depths, ranging from the surface to the deep chlorophyll maximum and the mesopelagic zone. Detection of Gp3 in the TARA1 dataset (which uses successive particle separation steps with filters of different sizes (49)) succeeded predominantly in the viral (<0.22um) and small prokaryotic (0.1-0.22um) size fractions, with few hits in the filters representing larger pro- and eukaryotic cells. TARA2 uses only a single filter, but the global distribution of Gp3 matches that of TARA1, and notably indicates the presence of Gp3 in marine environments from the Arctic to the Antarctic.

**Figure 2:**
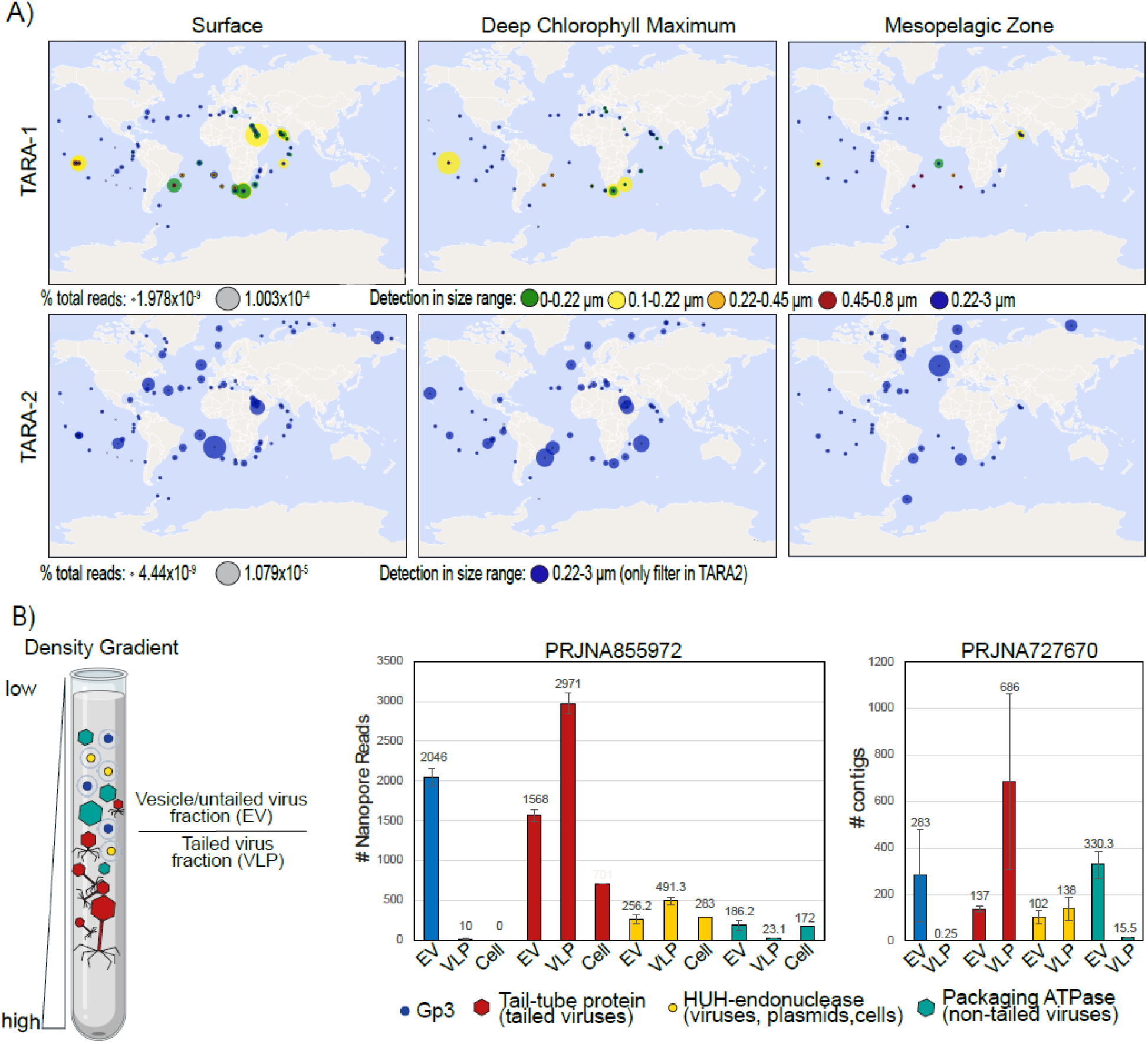
Global and local distribution of Gp3 in ocean metagenomes. A) Distribution and percentage of total reads mapping onto Gp3 in TARA Ocean 1 & 2 datasets at different depths and filter sizes, as assessed by Ocean Gene Atlas (46). B) Distribution of Gp3 and *Duplodnaviria* tail-tubes (VFAM00058), HUH-endonucleases (VFAM00122) and *Varidnaviria* packaging ATPases (VFAM00936) in two different density-gradient fractionated datasets. For PRJJNA855972, the EV fraction was defined as ranging from 1.14-1.19 g/ml in an iodixanol gradient, and for PRJNA727670 from 1.3-1.35 g/ml CsCl (4, 50, 51).

The maximum detected abundance of Gp3 was 0.0001% of all reads and 5 copies/cell. For comparison, a search for tailed phages using a tail-tube HMM VFAM00058 found a maximum of 0.0004% of reads and 470 copies per cell. Notably, no instances of Gp3 could be found in transcriptomes or associated with specific cellular metagenomes. Overall, this pattern of detection (absence in transcriptomes and presence in TARA ocean datasets which were prepared with methods precluding the detection of single-stranded DNA (52)) points towards the Gp3-carrying MGEs having double-stranded DNA genomes.

To investigate the occurrence of these MGEs on a finer scale, we focused on recent marine metagenomic studies where viral and cellular fractions had been separated first by size-exclusion, and viral fractions subsequently separated by density gradients (4, 50, 51). This resulted in a fraction defined as enriched in virus-like-particles (VLP, consisting of large viruses, particularly tailed *Duplodnaviria*) and a fraction enriched in both extracellular vesicles (EVs, which contain both cellular DNA and MGEs) and small, non-tailed viruses (*Monodnaviria*, *Singleaviria*, *Varidnaviria*). To confirm the quality of this separation, we first searched the fractions from both datasets for the presence of genes indicative of the presence of such viruses and MGEs. As expected, VLP, EV and cellular fractions were rich in HUH-endonucleases (VFAM00122) known to be found in virus particles, extracellular vesicles and cellular plasmids alike (53). Tail-tubes (VFAM00058) typical of tailed *Duplodnaviria* were detected predominantly in the VLP fractions while packaging ATPases (VFAM00936) typical of *Varidnaviria* were more abundant in the EV fraction. In contrast, Gp3 was detected almost exclusively in the EV fraction, with an average of less than 10 Gp3-positive nanopore reads or <1 Gp3-positive contig in the VLP fraction, and none in the cellular fraction. (Fig. 2B). As such, it appears that genomes encoding the Gp3 protein are found either encased in small virions or extracellular vesicles.

### Diversity and gene content of Charybdis elements

While some of the Gp3 encoding MAGs were circular, genomically cohesive and thus considered complete Charybdis elements, many others were shorter and fragmentary, leading to the possibility that Gp3 could also be present in a variety of other MGEs. With the intent of recovering high-quality, complete genomes, we next queried the IMG/VR v4 database of isolated and metagenomically detected viruses (54) for genomes encoding the Gp3 protein. We recovered 5819 hits in the IMG/VR database, 2,975 of them with direct terminal repeats and therefore presumably circular and complete **(Table S2)**.

The size of complete circular genomes ranges from 3,706 to 5,181bp, with a median size of around 4.6kb, similar to the 4.3 size median of the mediterranean sea dataset. Larger size outliers are present exclusively in the form of concatemers of multiple identical genomes, indicating that sequences encoding Gp3 are not simply fragmented parts of larger contigs (**Fig. S2**). All Gp3-positive IMG/VR genomes are derived from marine samples ranging from open ocean surfaces to sediments, including ocean trenches, hydrothermal vents and abyssopelagic zones (Fig. 3A). Further searches in the IMG/PR v1 database of plasmids (55) did not yield any results, indicating that MGEs encoding gp3 are exclusively found in marine environments and do not show any known hallmarks of plasmids.

**Figure 3:**
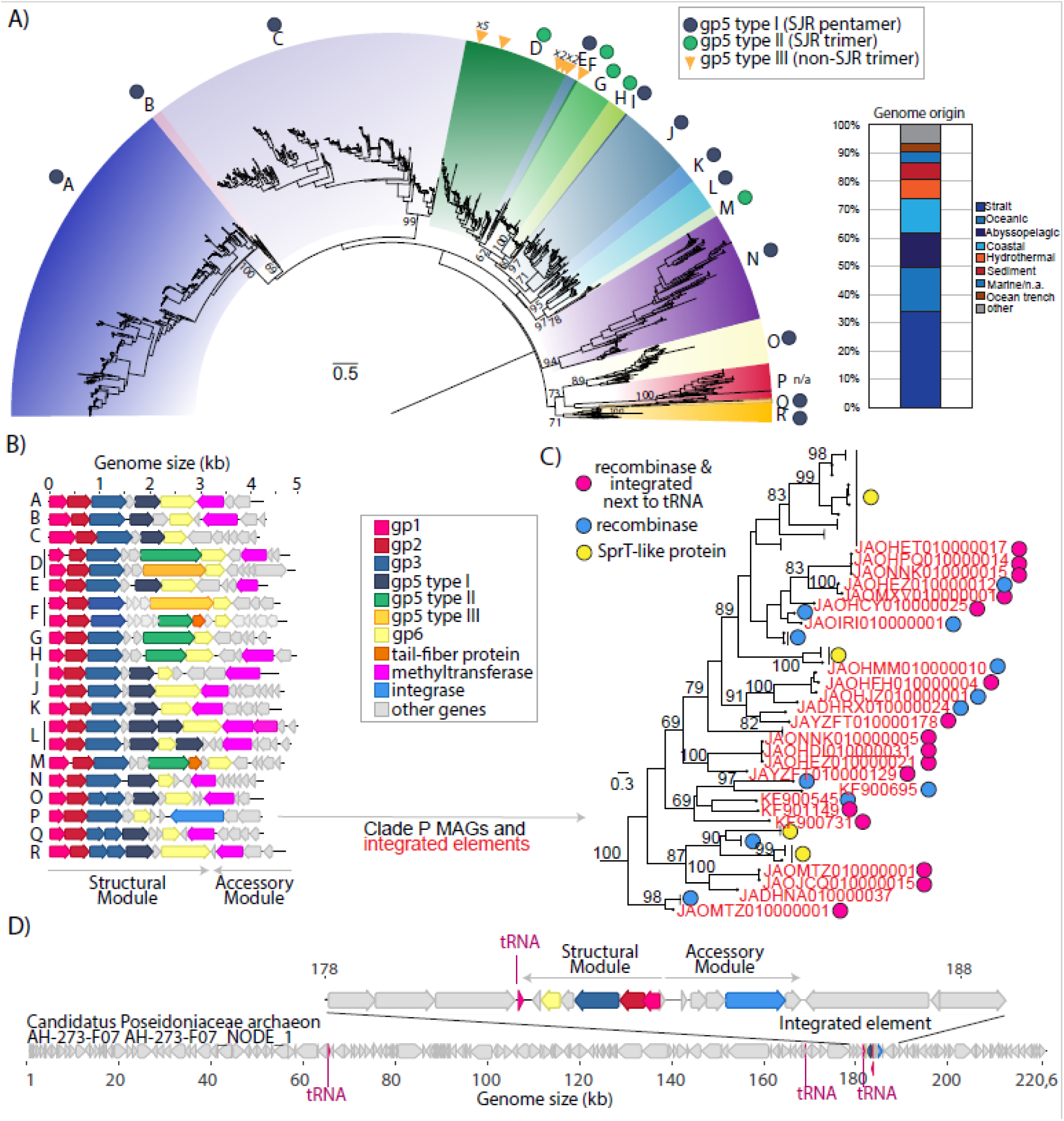
Diversity of Charybdis elements. A) Maximum likelihood phylogeny of Gp3 proteins found in complete circular IMG/VR v4 genomes. For genomes where instead of a single Gp3 protein, two genes encoded for four and three beta sheets separately (clades O,Q,R), amino acid sequences were concatenated and treated as one protein. Branch length indicated in legend corresponds to 0.5 aa substitutions/site. Relevant bootstrap values from 100 replicates are indicated on the tree. Bar graph indicates sample origin of both complete (circular) and non-complete (linear) genomes. B) Representative genomes from clades indicated in A. Circular genomes were arbitrarily linearized with gp1 as the 5’-end. C) Maximum likelihood phylogeny of Gp3 proteins of Clade P elements in A. NCBI Accession numbers of proviral MAGs are indicated in red. Clades of (nearly) identical Gp3 protein sequences from IMGVR were collapsed. D) Example of element integrated into genome of Candidatus Poseidoniaceae archaeon MAG. Tree rooted on clade O genomes (not shown).

A phylogenetic tree based on the conserved Gp3 protein (shared between all genomes with at least 30% amino acid identity) separates the Charybdis elements into multiple well-supported clades (Fig. 3a). These elements in IMGVR/V4 are separated into 2026 vOTUs of 95% average nucleotide identity over 85% of the genome (**Table S2**) – for comparison, 5709 genomes of the ubiquitous tailed phage family *Ackermannviridae* cluster into 3425 vOTUs in IMG/VRv4, indicating considerable sequence diversity within this group.

Regardless of clade, all genomes share the same basic structure as the initially described Charybdis element, indicating that Gp3 is indeed a hallmark for a novel, internally cohesive group of MGEs (Fig. 3B): Two gene clusters face different directions on the genome, varying in length depending on the genome. The first gene cluster universally encodes a group of three genes gp1-gp3 and an additional variable number of genes downstream of it. Interestingly, in three basal clades O, Q and R, the two Gp3 protein domains are split into two different genes. Notable variation is present among genes in the gp5 position, which encodes a SJR-protein in the prototypical Charybdis genome described in Fig. 1. These genes appear to encode virus-like protrusions which we will describe in more detail later, and which can be separated into three types: Most clades encode short Type I SJR-pentamers. Clades D, F, G, H and M encode longer Type II protrusions instead. Twelve instances of Type III non-SJR trimers are found sporadically in clades D and F, with their close relation to genomes encoding Type II SJR-trimers indicative of horizontal gene transfer events. Furthermore, clade P genomes lack any gp5 genes whatsoever, while some genomes within clade L and N encode multiple Type I SJR-pentamer genes. All clades additionally encode a hypothetical gene of unknown function downstream of gp5 (see gp6 in Fig. 1 and 4), predicted to form multiple alpha-helices of various lengths and number, connected by loop regions (See Fig. S1A).

**Figure 4:**
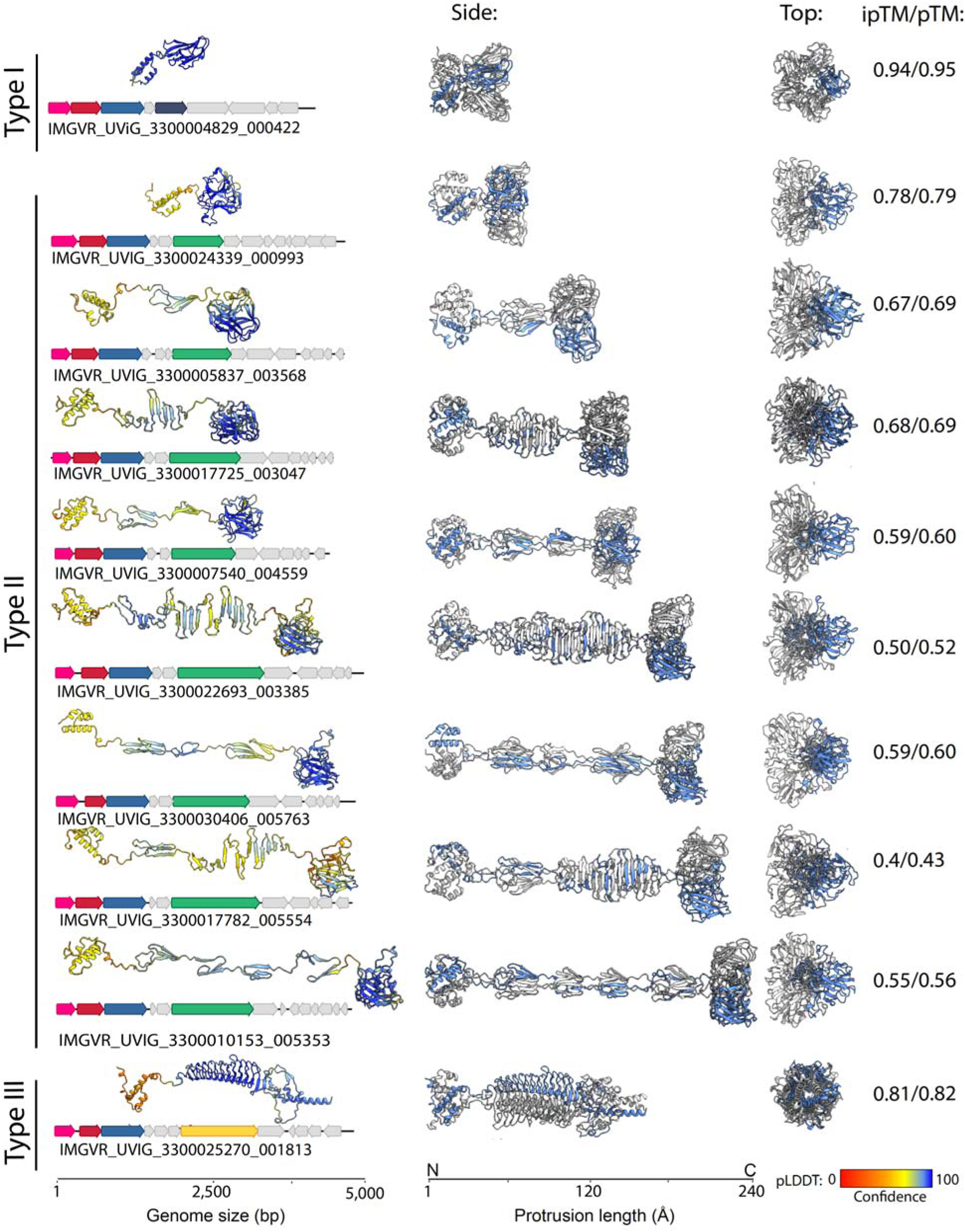
Assorted (predicted) protrusions formed by genes in the structural modules of Charybdis elements. Colored genes in the example genomes for each protrusion type correspond to gp1-3 (pink,red,blue) and gp5 (dark blue, green, yellow). Alphafold3 pTM score represents the predicted template modeling confidence, with > 0.5 suggesting that the predicted complex might be similar to the true structure. Alphafold3 ipTM scores represents the interface predicted template modeling confidence, measuring the confidence in the relative positions of the subunits in the multimer. pTM > 0.5 suggests that the predicted complex might be similar to the real structure. ipTM score 0.8 or above suggest high confidence predictions, below 0.6 suggest likely failed predictions. pLDDT (predicted local distance difference test represents a per-atom confidence score of predictions, with 100 (blue) representing the highest confidence and 0 (red) the lowest.

The oppositely oriented gene cluster is enriched in what appear to be accessory genes – in most instances one or more methyltransferases per genome, but also phosphoadenosine phosphosulfate reductase-like proteins, pyridoxal-phosphate epimerases, topoisomerases, integrases, MOM-like DNA modification proteins, SprT-protease like proteins and a large number of other hypothetical genes (**Fig. S3**) We thus term this operon the “accessory module” of the MGE, and the other gene cluster accordingly “structural module”. Save for topoisomerases, which occur in <100 genomes and exclusively in clade C and N, we could not detect any gene that can be reasonably interpreted as serving a role in replication of their circular genomes.

Only a single, incomplete Charybdis element from the IMG/VR database (IMGVR_UViG_3300020056_002831) could be assigned a host group (Marine group II archaea) based on k-mer matching, with the comprehensive database lacking both integrated elements and CRISPR-spacers targeting the genomes (Table S2). As mentioned before, homology searches against the NCBI database yield hits against metagenomically assembled contigs corresponding to a wide range of taxa (see Tables S1 and S3). However, closer inspection reveals that almost all hits are small contigs corresponding to just Charybis elements or their fragments, with very similar hits binned into genomes of taxonomically divergent bacteria (e.g. Pseudomonadota and Bacteroidota). Yet for 24 hits, we could confirm integration of Charybdis elements from Clade P (previously noted as lacking gp5-encoded putative protrusions) into the genomes of Marine Group II archaea, particularly within Candidate order Poseidoniales and the metagenomic type species Candidatus Poseidonia alphae (Fig. 3C **and D**) (38).

In contrast to circular Charybdis elements, these proelements are integrated into their host in such a way as to orient the structural and accessory modules facing away from each other. Additionally, several elements are integrated next to host tRNA genes, which is a typical insertion site of mobile genetic elements (56). As all integrated elements (but not all members of the P clade) contain a gene annotated as integrase/recombinase, it is likely that the acquisition of such accessory genes has allowed some clade P Charybdis elements to target tRNAs of marine group II archaea for integration.

### A variable genomic locus is predicted to encode protrusion-like structures resembling viral receptor binding proteins

Curiously, 58% of the Charybdis elements in IMG/VR v4 are taxonomically annotated as members of the tailed prokaryotic virus class *Caudoviricetes*. Taxonomic assignment using vContact3 (https://bitbucket.org/MAVERICLab/vcontact3/src/master/) places the elements into a new order within the *Caudoviricetes*. However, the elements form a clearly separate cluster of genomes with no proteins shared with other viruses (**Fig. S4**). Additionally, Charybdis elements are significantly smaller than even the smallest predicted members (7.4 kb) of the *Caudoviricetes* (33). Further, they do not encode the characteristic HK97-fold hallmark capsid protein or any other gene for proteins typical of in either tailed phages or known phage satellites, such as terminases, portal proteins or prohead proteases (**Fig. S1a**) (57). Upon closer inspection, the taxonomic assignment appears derived from the annotation of certain genes in the gp5 position as associated with viral tail structures. These genes are characteristic, but not diagnostic of *Caudoviricetes* genomes. Indeed, structural predictions yielded multimeric assemblies with similarities to the protrusions that adorn various viral capsids (Fig. 4, Fig. 5A)

**Figure 5:**
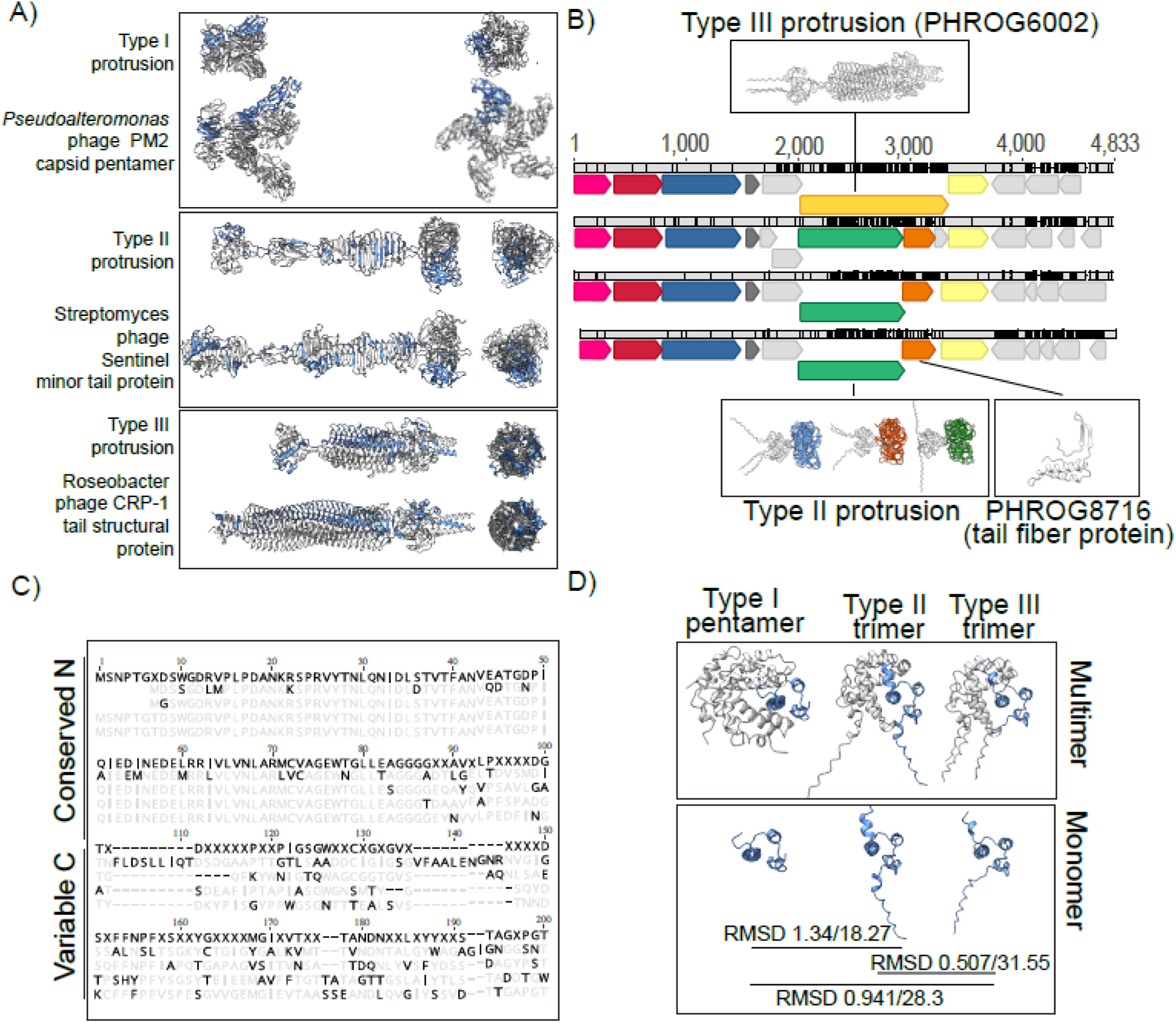
Resemblance, recombination and modularity of putative protrusion proteins. A) Comparison of three protrusion types with known protrusions of other viral elements. Charybdis protrusions, Streptomyces phage sentinel Uniprot ID A0A873WQA2 and Roseobacter phage CRP-7 uniprot ID A0A646QVI9, were predicted using Alphafold3, PM2 capsid pentamer is derived from PBD ID 2W0C. B) Genome alignment of four closely related genomes. Arrows represent genes, grey bars aligned genomes (IMGVR_UViG_3300001524_000321, IMGVR_UViG_3300028892, IMGVR_UViG_3300001680_000868 and IMGVR_UViG_3300022693_002222, respectively), with black notches indicating single nucleotide polymorphisms. C) Alignment of first 200 amino acids from protrusions in B. Topmost sequence in dark letters represents consensus, greyed out letters represent agreements, dark letters disagreements to consensus. Order of sequences is the same as in B. D) Conserved structures of N-terminal knob domain of type I (IMGVR_UViG_3300004829_000422), II (IMGVR_UViG_3300017782_005554) and III (IMGVR_UViG_3300025270_001813) protrusions. Root mean square distance (RMSD) for N-terminal knob and entire structure are indicated as a measure of structural overlap between monomers.

The hypothetical gene products of gp5 can be roughly grouped into three types based on predicted structures: Type I proteins average 170 aa in length, with an N-terminal anti-parallel triple-helix bundle and a C-terminal single jelly-roll fold-like domain, connected by a short loop region. DALI searches shows homology of the SJR domain to pentameric nucleoplasmin-like single jelly-roll folds (DALI Z-score 9.5 to pentameric histone deacetylase HDT (pdb id 7VMI). In accordance with this, proteins from this type are confidently predicted (iPTM score 0.94) to form pentamers with a ring-like alpha helical structure at the N-terminus, from which the five SJR domains are angled outwards in the appearance of a flower. This notably differentiates their structure from the nucleoplasmin-fold proteins that form inward-angled pentamers on the capsid such as that of satellite tobacco mosaic virus (pdb id: 7M3R). The predicted structures more closely resemble (but display no apparent homology to the pentameric decorations on the capsids of *Varidnaviria* phages such as *Pseudoalteromonas* phage PM2 (Fig. 5A). We also performed alternative multimeric assemblies as trimers, however, these yielded incomplete pentamers (missing two proteins) and considerably lower prediction confidence scores (**Fig. S5**).

Type II proteins also contain and N-terminal anti-parallel triple-helix bundle, but continue into a “stalk” of variable length. As expected, overall prediction confidence scores decrease for longer stalks, dipping from 0.78 for the shortest to 0.5 for the longest protein. While the stalks are predicted to form different beta-sheet “rib” and “bulb” elements, all protrusions end in a C-terminal SJR fold domain of the lectin-type (DALI Z-score 15.6 to the trimeric Z-protein at the 3-fold axis of symmetry in *Anabaena* phage A4L capsids [*Duplodnaviria*] (58)). In accordance to this, and unlike type I proteins, the higher confidence multimeric prediction for those proteins is that of a trimer (ipTM 0.78 vs 0. 29 for the shortest protrusion as trimer and pentamer, respectively, **Fig. S5**). The lectin-type SJR trimer attached to a stalk of various length overall gives an appearance similar to the tail fibers of *Duplodnaviria* (Fig. 5a). Hidden Markov model hits against several Prokaryotic Virus Remote Homologous Groups database ((59) PHROGS 15,388; 21,017; 29,228) are all annotated as unknown functions, but similar PHROGS (e-value <0.05) are annotated as tail fiber proteins, structural proteins or (minor) tail protein. Additionally, genes downstream of some of these genes generate hits to PHROGS groups 4912 and 8716, which are annotated as tail fiber/virion structural proteins Type III proteins are the largest (∼400aa) and rarest (found only in 12 genomes), with an entirely different predicted structure, save for the N-terminal anti-parallel triple-helix bundle. They lack SJR-domains but display the characteristic appearance of trimeric viral tail-spikes, to which their sequences are indeed homologous (PHROGS 4636, 6002) (Fig. 5A). Again, alternative assemblies as pentamers are considerably worse (**Fig. S5**).

While structural homology exists between both type II and III protrusion proteins and proteins of known viruses, sequence-based searches only yield high-similarity hits to either short, dubiously binned metagenomic contigs (Charybdis elements themselves) or low coverage hits with low amino acid identity – as such, we cannot ascertain whether Charybdis element encoded protrusion genes were acquired from a specific cellular or viral donor (**Table S3**). However, comparison of the available genomes provides evidence of horizontal gene transfer leading to the exchange of these putative protrusions between Charybdis elements themselves (Fig. 5B). In analyzing a group of four closely related genomes, three genomes encode short Type II proteins. While the 5’-end of the gene (corresponding to the alpha-helical domain) is highly conserved (>90% ANI), the 3’-end, (encoding the SJR-domain), is highly variable (<30% ANI). A fourth, largely identical genome encodes a Type III protein, however, its N-terminus shows a high degree of sequence identity to the N-termini of Type II proteins from the other genomes (Fig. 5C). Indeed, it appears that while the sequence of the 5’-end of protrusion genes is not necessarily conserved between all three types of protrusion-forming proteins, the structures of their N-terminal anti-parallel triple-helix bundle structures largely overlap (highest root mean square distance < 1.4 angstrom for the N-terminus vs lowest root mean square distance 18.27 angstrom for the overall structures) (Fig. 5D). Overall, the conserved N-terminal and variable C-terminal structures that appear to undergo recombination indicates a modular layout typically seen in proteins involved in interactions with cellular hosts.

## Discussion

We have discovered a novel, widespread, genomically cohesive group of marine mobile genetic elements that we term Charybdis elements. Due to the metagenome-only nature of our analysis and the lack of clear homologies to known mobile genetic elements including viruses, we can at this point only speculate on what type of mobile genetic element their represent (Figure 6.)

**Figure 6:**
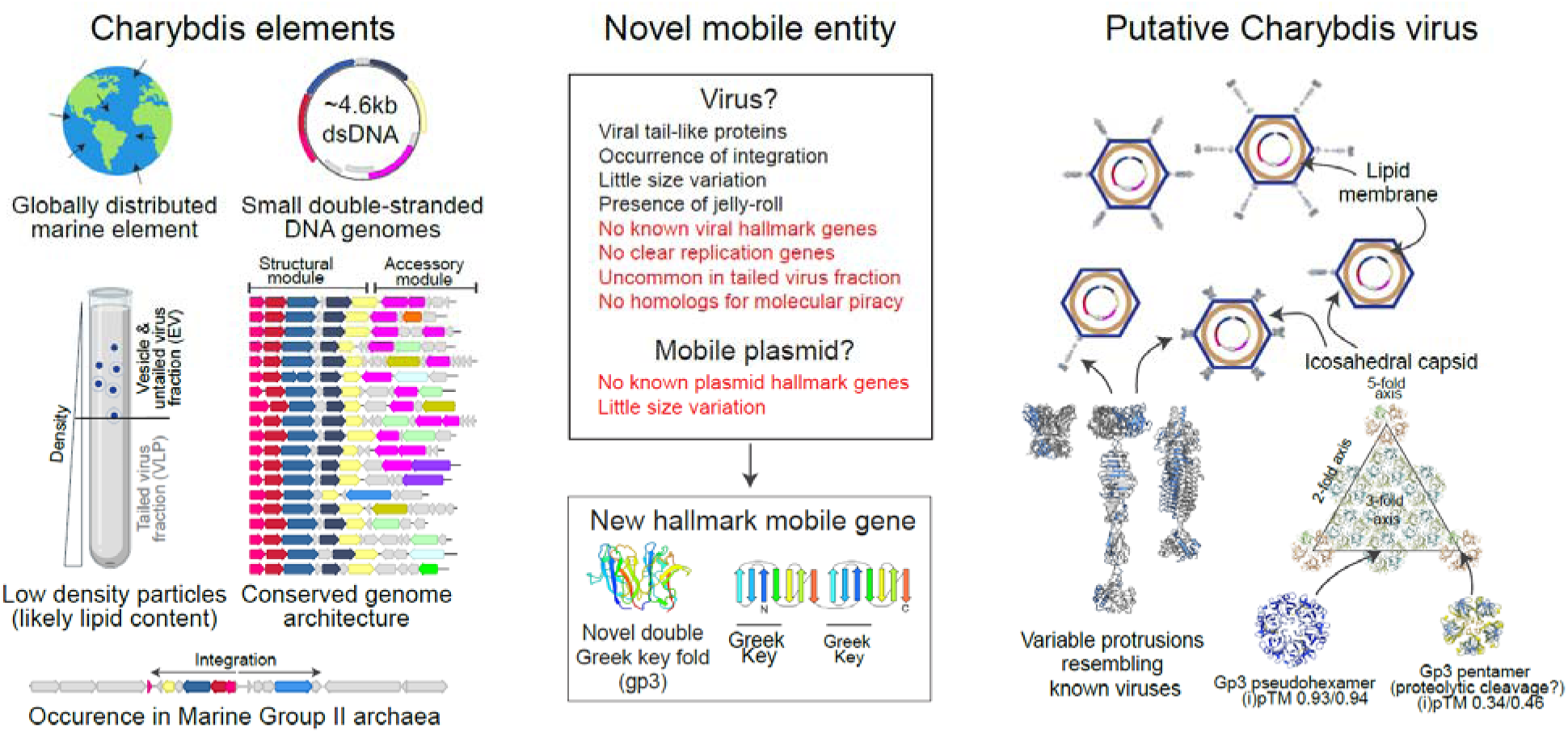
Summary of evidence supporting and refuting various hypotheses on the nature of Charybdis elements. Note that the T=12 capsid structure in the upper right corner was arbitrarily chosen.

Charybdis genomes are circular, uniformly around 4.6kb in size and, based on their detection in metagenomes that exclude RNA and single-stranded DNA, appear to have double-stranded DNA genomes. Their hosts, which we predicted based on occasional genomic integration events, are marine group II archaea which have recently been combined into the Candidate Order Poseidoniales(38). These uncultured but extremely common photoheterotrophic archaea display global marine distribution in a wide range of oceanic depths, overlapping with that of Charybdis elements (60).

Charybdis elements encode a number of accessory genes with potential roles in enabling evasion of their host’s defenses, manipulation of metabolic pathways and (rarely) integration into their chromosomes – a common characteristic of plasmids, transposons and virus-like elements. Plasmids and other vesicle-borne MGEs can greatly expand and contract their genome size to accommodate additional accessory genes [for example (61, 62)] – an observation which we notably have not been able to make, despite the availability of thousands of Charybdis elements in a variety of marine environments. Furthermore, we note a decisive lack of plasmid-associated genes, which also speaks against these elements being plasmids. Similarly, Charybdis elements share no similarities to recently described EV-encased transposable elements (3, 50), although they are abundant in the datasets provided in these studies (see Fig. 2B). Regardless, these MGEs could represent a novel group of marine elements encased in the lipid vesicles of their hosts. The protrusions encoded in their genomes could function as cellular attachment factors, conferring infected hosts with the ability to bind to novel surfaces (examples of such phage-tail like structures exist in bacteria such as Bdellovibrio (63).

However, since these ubiquitous protrusions are predicted to form structures resembling those of, or even being homologous to, virus-like protrusions, the most likely (but by no means definitive) hypothesis is that these elements represent a hitherto unknown form of virus-like or associated element. We did not detect any known hallmark proteins, hence Charybdis elements cannot clearly be classified as members of an extant viral realm. Some lineages of the Charybdis elements encode genes homologous to parts of *Caudoviricetes* tails. However, they are missing most genes required to build these elaborate structures – assuming that these proteins do not form much simpler protrusions, this indicates that they could be satellite MGEs associated with these viruses. Numerous satellites appropriate virus structural and replicative elements to package multiple copies of their small (but still generally 10kb+) genomes into the heads of tailed phages (57). However, within virus enriched datasets, Charybdis elements are almost completely absent from the large tailed virus fractions – as such, it does not appear as if multiples of their small genomes were packaged into normal-sized tailed virus capsids. Enriched in low-density fractions, their genomes could be encased in smaller satellite virus particles, providing elements of their tail themselves and pirating capsid and other structural elements from ubiquitous *Caudoviricetes* viruses. Their likely host, the Poseidoniales archaea, are indeed infected by tailed viruses that could provide the missing elements of the Charybdis replication cycle (64, 65). Supporting their nature as satellites, we could not identify a gene encoding a known replication associated protein – barring one or more novel proteins facilitating replication, these elements likely depend on other replicators. The occasional detection of genome concatemers indicates that the elements replicate using a rolling-circle replication mechanism, and genes enabling this could indeed be provided by helper *Caudoviricetes* (66). Alternatively, their genomes could be segmented or multipartite and encode replication associated proteins on a different molecule that we failed to identify. However, such genome types rarely or never infect prokaryotes(67), and to date no double-stranded DNA elements with segmented or multipartite genomes are known. Additionally, no segmented/multipartite MGES are known to integrate into cellular chromosomes, making it unlikely that these elements are indeed segmented/multipartite. The absence of clear replication associated genes in viruses is however not unprecedented – for example, some members of the archaeal *Pleolipoviridae* apparently replicate using host factors only (68). What speaks against these elements being satellites is that all known *Caudoviricetes* satellites encode genes homologous to their virus targets, allowing them to reorganize virus assembly to facilitate the packaging of their own genomes (57) – such genes are entirely absent in Charybdis elements, which of course does not rule out that some genes might nonetheless assume such functions. Finally, a large, phylogenetically distinct number of Charybdis elements do not encode trimeric protrusions typical of *Caudoviricetes* – instead, they appear to encode genes for pentameric protrusions more expected on members of *Varidnaviria*.

Alternatively, Charybdis elements could form small virus capsids akin to the *Microviridae* or *Varidnaviria* by themselves, or they could be contained within vesicles similar to archaeal virus-like plasmid pRISE or *Pleolipoviridae*. The uniformity in genome size supports the former assertion: constraints of small virus capsids do not allow for significant size variation. Barring an envelope-only virus, which are rare among prokaryotes, the most likely candidate for a capsid protein encoded in the genome (rather than appropriated from a helper virus) is the double Greek-key fold forming Gp3. Keeping in mind the highly speculative nature of this thought, Gp3 superficially resembles double-jelly-roll fold (DJR) proteins of the *Varidnaviria*. The capsid faces of *Varidnaviria* (e.g. PM2, PDB id 2w0c) are formed by trimers (pseudohexamers) of DJR-proteins and indeed, high-confidence multimer prediction of the Gp3 protein (ipTM 0.93) shows a remarkable similarity to *Varidnaviria* pseudohexamers (Fig. 6).

In *Varidnaviria*, dedicated SJR proteins form pentamers at the twelve vertices of their icosahedral capsids. These SJR pentamers are often decorated with trimeric tail fibers (see for example the iconic adenovirus). SJR-proteins are found in almost all Charybdis elements, however, they appear to be part of longer protrusions more likely to attach to vertices than to form them themselves. Hypothetically, the vertices could be formed by single Greek-key fold Gp3 variants, derived perhaps from proteolytic cleavage at the linker between the two parts of Gp3. We also note that the basal clade O, Q and R genomes, instead of encoding a Gp3 protein with two beta-sandwich domains, encode two successive single beta-sandwich proteins, which are predicted to assemble both as hexamers and pentamers (albeit the latter with much lower confidence, see Fig. 6). With the potential to form both pseudohexamers and pentamers covering the face of a capsid, it is possible to imagine virions resembling those of many members of the *Varidnaviria*, perhaps containing an internal or external membrane, and sporting either one or multiple protrusions mediating attachment to their hosts (Fig. 6)

In our structural phylogeny, Gp3 proteins cluster outside of all known viral or cellular DJR proteins and at least visually resemble vertebrate gamma-crystallin proteins. (Gamma) crystallins are a group of proteins involved in a wide variety of roles in both pro- and eukaryotes (69). Notable crystallins form lens proteins of the vertebrate eye, heat shock proteins, and a number of enzymes. So far, no crystallins have been implicated in a structural role for virus capsids. As most viral capsid proteins are descendants of cellular proteins, this could hint at the presence of a novel viral realm (70). Given that their only known hosts are widespread but unculturable marine archaea, the determination of the true nature of these elements is not possible at this point. Regardless, one can at least imagine (or perhaps hallucinate) a virus strongly resembling a small member of the *Varidnaviria* in shape, if not genome content or size, that has escaped detection until now.

## Supporting information

Supplemental Datafile 1

## Supplementary Data

Supplementary datafile 1: Curated complete, circular Charybdis genomes analyzed in this study

Supplementary Table S1: Homology search results for all genes of genome depicted in Figures 1, S1

Supplementary Table S2: List of all Charybdis genomes in IMG/VR v4 Supplementary Table S3: Homology search results of exemplary gp5 protrusion types

## Acknowledgements & Funding

The authors would like to thank Simon Roux, Kathryn Kauffman and Reed Stubbendieck for their valuable input. P.K. is funded by N.I.H. Grant 5P20GM152333-02, A.L. is funded by N.S.F. grant 2428961.

## Author Contributions

P.K. planned and executed the study and performed analyses. A.L. contributed crucial information on possible capsid geometries. P.K. and A.L. wrote the manuscript.

## Data availability Statement

The authors confirm that all the data used in this manuscript (accession numbers, genomes) are available in the manuscript, its supplementary files or upon request from the authors.

## Methods

### Detection of Charybdis elements from (meta)genomic datasets

SRA files from Bioproject PRJEB30684 were downloaded and unpacked using the NCBI SRAtoolkit 3.2.1 (https://github.com/ncbi/sra-tools) and reads were trimmed and quality-controlled using fastp (71), standard settings. Reads were then assembled into contigs using MEGAHIT 1.2.9 (72), standard settings. All contigs between 1 and 15kb were then annotated using PHAROKKA 1.5.1 (73), metagenomic settings. Since our initial search targeted microviruses, network graphs and clustering into groups of closely related genomes was performed using MOP-UP (35), standard settings except using an amino acid identity cutoff of 30% to cluster proteins. Resulting subgroups of genomes in the Master.csv file were sorted by abundance and manually inspected for genes of interest through visualization in Geneious Prime (Dotmatics); blastp (74)/skyblast (https://sky-blast.com) searches against the NCBI nr database; Foldseek(42) searches against all available databases in July 2025, Alphafold3 (75) webserver reconstructions and DALI searches of predicted structures against the PDB database (44).

Further research focused on the third-most abundant (by number of genomes) group in ERR3063491, in which all genomes contained the gp3 gene that was visually identified using ChimeraX1.8 (76) to encode an interesting double Greek-key fold protein. All gp3 genes were extracted, translated and aligned using Clustal Omega 1.2.2 (77) with three refinement iterations, evaluating the full distance matrix, using the Geneious Prime GUI. The untrimmed alignment was then used to create a hidden Markov model (hmm) using HMMER 3.4 (78).

Subsequently, HMMER hmmsearch on all proteins from Bioproject PRJEB30684 contigs were performed using the aforementioned gp3 hmm. Genomes with hits of e-value < 0.001 were extracted and manually inspected for false positive hits using Geneious Prime and Alphafold3 reconstructions, resulting in a list of 710 confirmed gp3 positive genomes. The same procedure to detect Gp3 was performed for PRJNA727670 and PRJNA855972. Contigs from PRJNA727670 were assembled as described above. Nanopore reads of PRJNA855972 were downloaded from https://zenodo.org/records/11551382. The Gp3 hmm was also used as input for query of all datasets contained in the Ocean Gene Atlas v2.0 (46) at https://tara-oceans.mio.osupytheas.fr/ocean-gene-atlas/ using the standard e-value threshold of 1E-10.

### Genome curation of IMG/VR v4 Charybdis elements

The aforementioned Gp3 hmm was used to query the IMG/VR v4 and IMG/PR v1 protein dataset of viruses and plasmids, and the list of hits was used to extract all gp3 positive hits from the IMGVR v4 contig dataset. We subsequently focused on genomes with direct terminal repeats, which we assumed to be circular and complete. We detected 100bp+ direct terminal repeats using the find repeats function in Geneious Prime and subsequently deleted one instance of each to obtain full genomes, which we subsequently annotated with PHAROKKA 1.5.1. All proteins were extracted and clustered at 30% amino acid identity using MOP-UP. Large clusters of proteins were retrieved, and hidden Markov models were generated for proteins Gp1, Gp2, Gp5 and Gp6. We then used these hmms to supplement the hmms contained in the PHAROKKA 1.5.1 database to re-annotate all genomes. We subsequently performed multiple rounds of manual refinement of annotation and gene calling by comparing closely related genomes for missed annotations, using visualization in Geneious Prime (Dotmatics); blastp (74)/skyblast (https://sky-blast.com) searches against the NCBI nr database; Foldseek(42) searches against all available databases in July 2025, Alphafold3 (75) webserver reconstructions and DALI searches of predicted structures against the PDB database (44).

### Alignments and phylogenetic analysis

Phylogenetic trees of aligned Gp3 proteins were based on ClustalOmega (see above) alignments of gp3s from curated IMG/VR v4 genomes. For genomes where instead of a single Gp3 protein, two genes encoded for four and three beta sheets separately (clades O,Q,R), amino acid sequences were concatenated and treated as one protein. Maximum likelihood phylogenetic trees were constructed with 100 fast bootstrap replicates using RAxML-NG (79) using the GAMMA+WAG model. For structural phylogenetic analysis, proteins without resolved structures were predicted using Alphafold3 or downloaded from PDB. These proteins were subsequently used as input for the Foldtree (45) collab notebook at https://github.com/DessimozLab/fold_tree.

---

**Figure S1:**
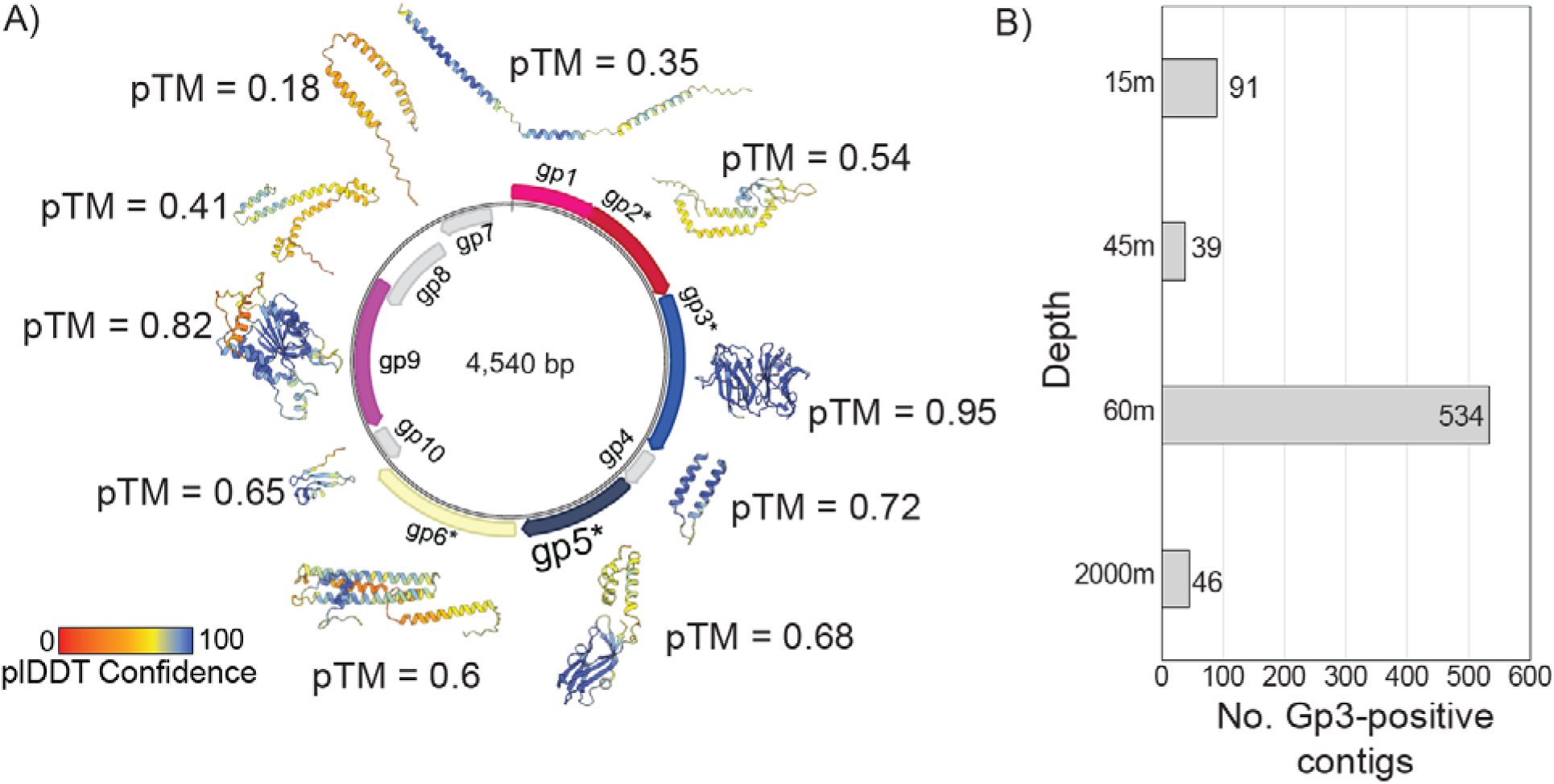
Structural predictions and depth distribution for Gp3-positive MAGs. **a)** Alphafold3 structure predictions of proteins encoded in an exemplary MAG. Arrows on circular genome indicate predicted genes, with asterisks indicating clear gene conservation across multiple genomes. Alphafold3 pTM score represents the predicted template modeling confidence, with > 0.5 suggesting that the predicted complex might be similar to the true structure. pLDDT (predicted local distance difference test) represents a per-atom confidence score of predictions, with 100 (blue) representing the highest confidence and 0 (red) the lowest. B) Number of contigs encoding Gp3 in mediterranean sea virome PRJEB30684.

**Figure S2:**
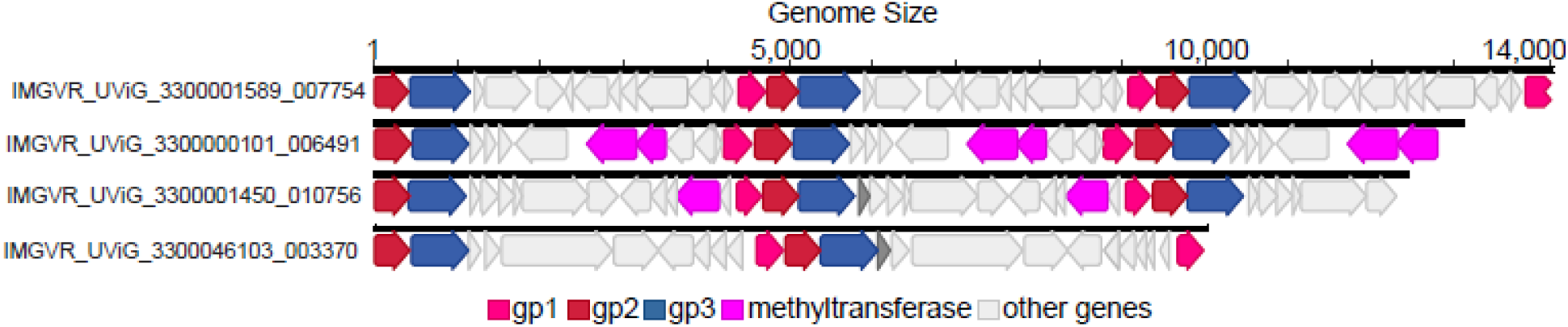
Examples of Charybdis genome concatemers in the IMGVRv4 database. Arrows indicate genes, gp2 was chosen as an arbitrary 5’ end of the genomes.

**Figure S3:**
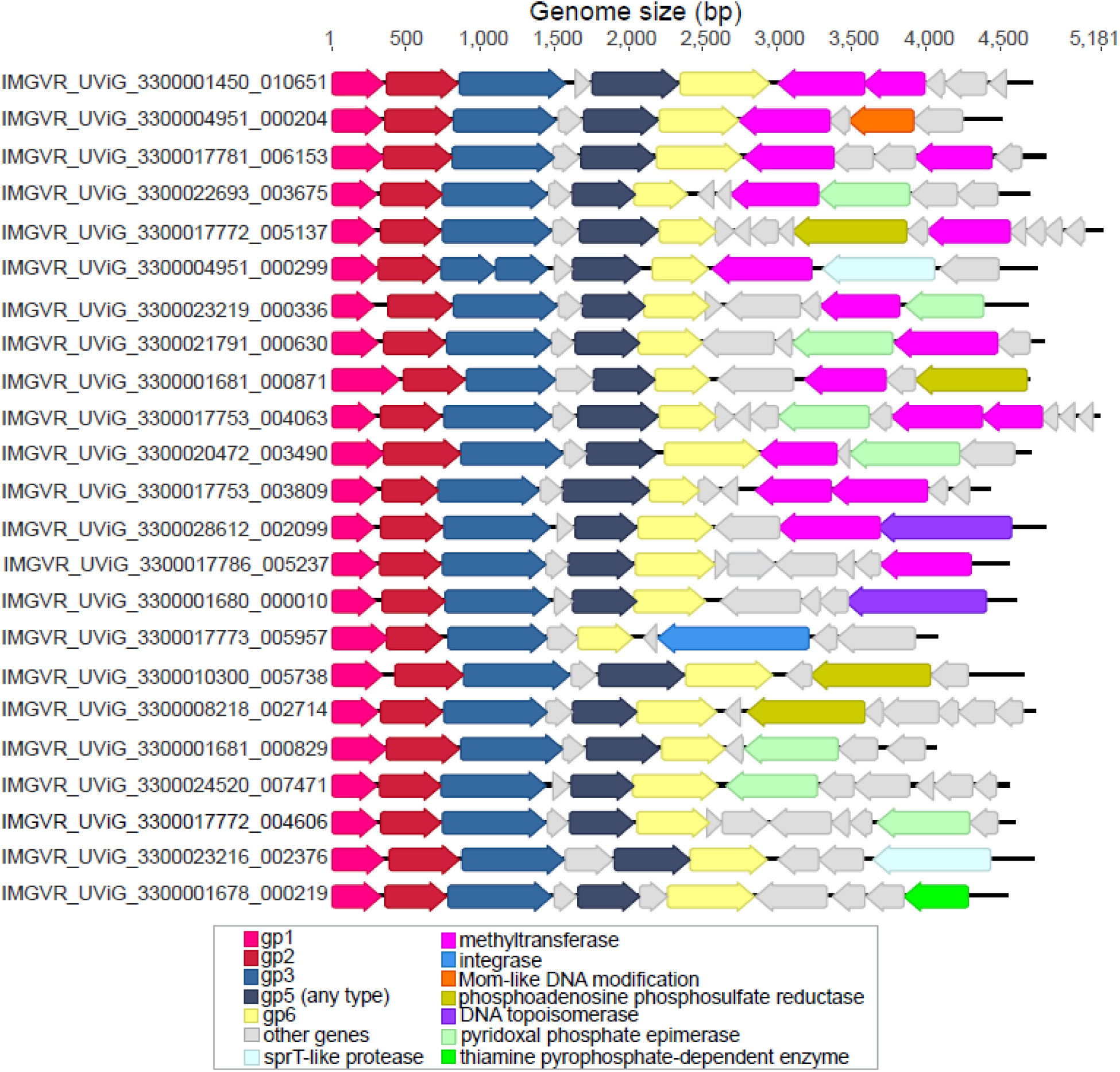
Gene content of select Charybdis elements, with focus on accessory genes. Arrows indicate genes. Circular genomes were arbitrarily linearized with gp1 as the 5’-end.

**Figure S4:**
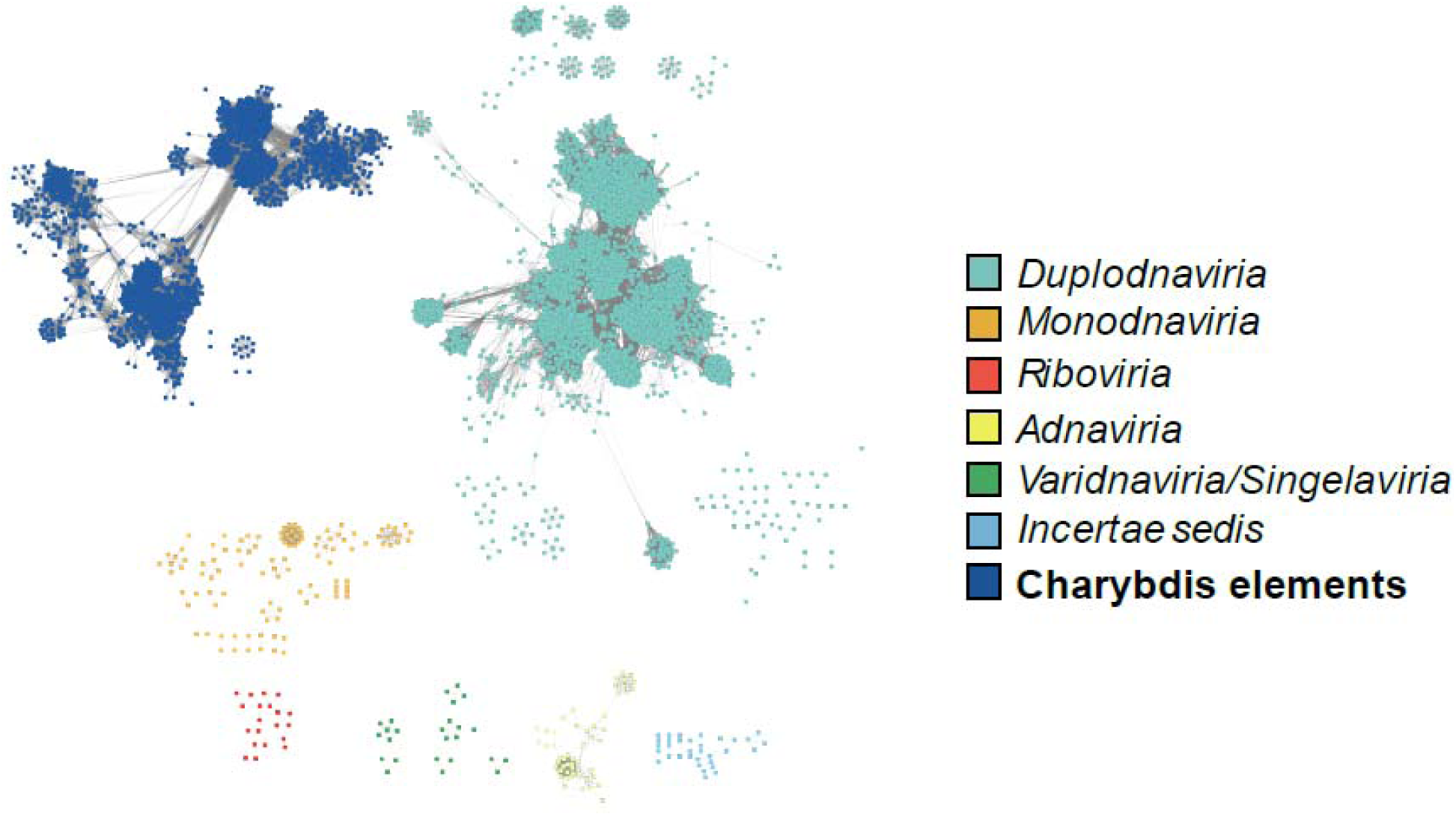
Protein sharing network of archaeal and bacterial viruses. Nodes represent individual genomes, connected by edges indicating shared proteins as calculated by vContact3 https://bitbucket.org/MAVERICLab/vcontact3/src/master/. Rectangles are coloured based on ICTV taxonomy.

**Figure S5:**
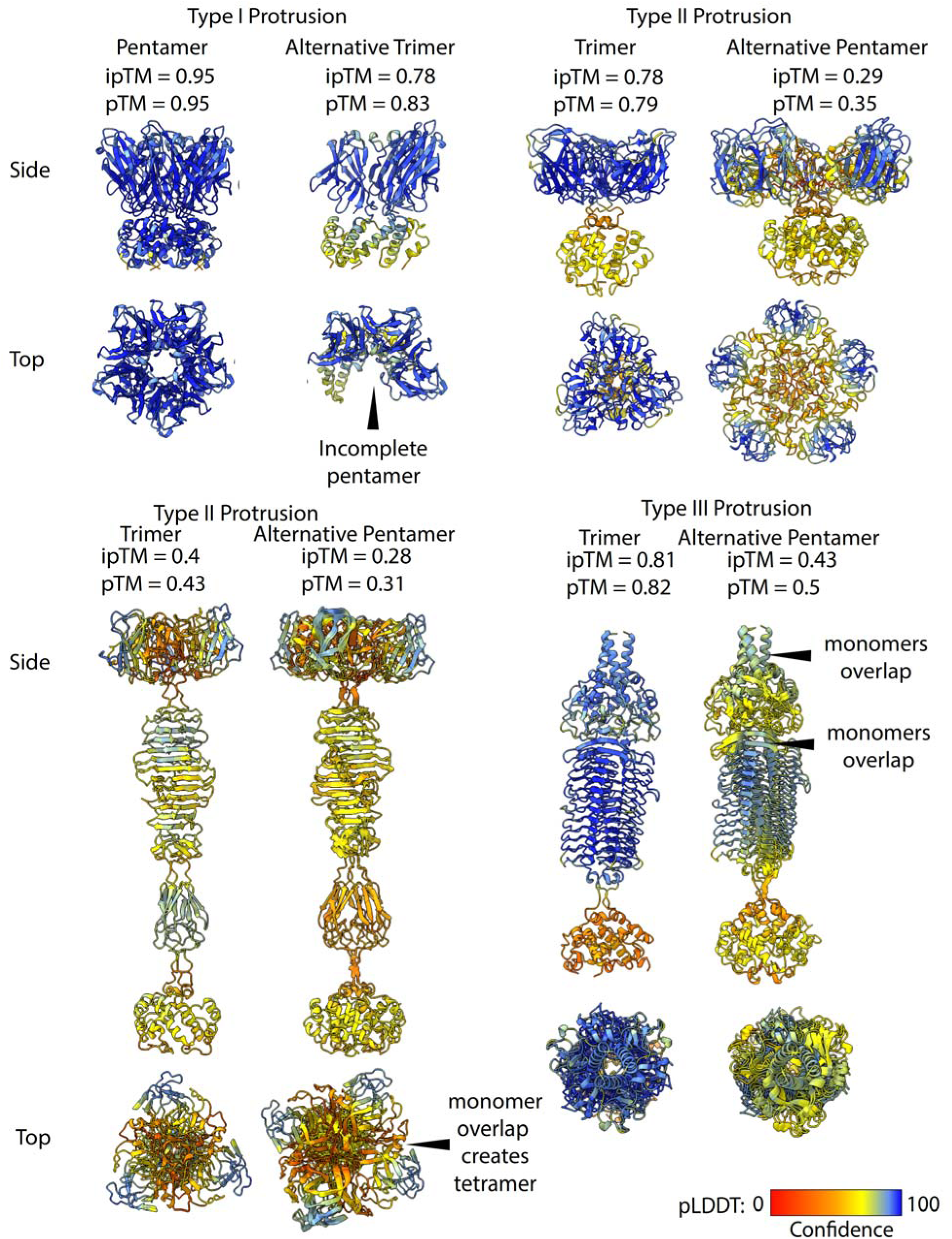
Alternative structural predictions and issues with Alphafold assemblies. Predictions are shown for IMGVR_UViG_3300004829_000422 (type I), IMGVR_UViG_3300024339_000993 and IMGVR_UViG_3300017782_005554 (type II) and (type III) IMGVR_UViG_3300025270_001813. pTM > 0.5 suggests that the predicted complex might be similar to the real structure. ipTM score 0.8 or above suggest high confidence predictions, below 0.6 suggest likely failed predictions. pLDDT (predicted local distance difference test represents a per-atom confidence score of predictions, with 100 (blue) representing the highest confidence and 0 (red) the lowest.

## References

1. Koonin EV, Dolja VV, Krupovic M, Varsani A, Wolf YI, Yutin N, Zerbini FM, Kuhn JH. 2020. Global organization and proposed megataxonomy of the virus world. Microbiol Mol Biol Rev 84:1–32.

2. Varsani A, Abd-Alla AM, Arnberg N, Bateman KS, Benkő M, Bézier A, Biagini P, Bojko J, Butkovic A, Canuti M. 2025. Summary of taxonomy changes ratified by the International Committee on Taxonomy of Viruses (ICTV) from the Animal DNA Viruses and Retroviruses Subcommittee, 2025. Journal of General Virology 106:002113.

3. Hackl T, Laurenceau R, Ankenbrand MJ, Bliem C, Cariani Z, Thomas E, Dooley KD, Arellano AA, Hogle SL, Berube P, Leventhal GE, Luo E, Eppley JM, Zayed AA, Beaulaurier J, Stepanauskas R, Sullivan MB, DeLong EF, Biller SJ, Chisholm SW. 2023. Novel integrative elements and genomic plasticity in ocean ecosystems. Cell 186:47–62 e16.

4. Eppley JM, Biller SJ, Luo E, Burger A, DeLong EF. 2022. Marine viral particles reveal an expansive repertoire of phage-parasitizing mobile elements. Proc Natl Acad Sci U S A 119:e2212722119.

5. Alqurainy N, Miguel-Romero L, de Sousa JM, Chen J, Rocha EP, Fillol-Salom A, Penades JR. 2023. A widespread family of phage-inducible chromosomal islands only steals bacteriophage tails to spread in nature. Cell host & microbe 31:69–82. e5.

6. Zheludev IN, Edgar RC, Lopez-Galiano MJ, de la Pena M, Babaian A, Bhatt AS, Fire AZ. 2024. Viroid-like colonists of human microbiomes. Cell 187:6521–6536 e18.

7. Koonin EV, Dolja VV, Krupovic M, Kuhn JH. 2021. Viruses Defined by the Position of the Virosphere within the Replicator Space. Microbiol Mol Biol Rev 85:e0019320.

8. Krupovic M, Dolja VV, Koonin EV. 2023. The virome of the last eukaryotic common ancestor and eukaryogenesis. Nat Microbiol 8:1008–1017.

9. Liu Y, Demina TA, Roux S, Aiewsakun P, Kazlauskas D, Simmonds P, Prangishvili D, Oksanen HM, Krupovic M. 2021. Diversity, taxonomy, and evolution of archaeal viruses of the class *Caudoviricetes*. PLoS Biol 19:1–26.

10. Krupovic M, Makarova KS, Koonin EV. 2022. Cellular homologs of the double jelly-roll major capsid proteins clarify the origins of an ancient virus kingdom. Proc Natl Acad Sci U S A 119.

11. Krupovic M, Kuhn JH, Wang F, Baquero DP, Dolja VV, Egelman EH, Prangishvili D, Koonin EV. 2021. Adnaviria: a New Realm for Archaeal Filamentous Viruses with Linear A-Form Double-Stranded DNA Genomes. J Virol 95:e0067321.

12. Pietilä MK, Roine E, Sencilo A, Bamford DH, Oksanen HM. 2016. Pleolipoviridae, a newly proposed family comprising archaeal pleomorphic viruses with single-stranded or double-stranded DNA genomes. Archives of virology 161:249–256.

13. Prangishvili D, Krupovic M, Consortium IR. 2018. ICTV virus taxonomy profile: Ampullaviridae. Journal of General Virology 99:288–289.

14. Krupovic M, Quemin ER, Bamford DH, Forterre P, Prangishvili D. 2014. Unification of the globally distributed spindle-shaped viruses of the Archaea. Journal of Virology 88:2354–2358.

15. Prangishvili D, Mochizuki T, Liu Y, Krupovic M, Consortium IR. 2019. ICTV virus taxonomy profile: Clavaviridae. Journal of General Virology 100:1267–1268.

16. Prangishvili D, Krupovic M, Consortium IR. 2018. ICTV virus taxonomy profile: Globuloviridae. Journal of General Virology 99:1357–1358.

17. Prangishvili D, Mochizuki T, Krupovic M, Consortium IR. 2018. ICTV virus taxonomy profile: Guttaviridae. Journal of General Virology 99:290–291.

18. Kim J-G, Gazi KS, Krupovic M, Rhee S-K, Consortium IR. 2021. ICTV virus taxonomy profile: thaspiviridae 2021. Journal of General Virology 102:001631.

19. Laso-Pérez R, Wu F, Crémière A, Speth DR, Magyar JS, Zhao K, Krupovic M, Orphan VJ. 2023. Evolutionary diversification of methanotrophic ANME-1 archaea and their expansive virome. Nature microbiology 8:231–245.

20. Huang L, Wang H, Consortium IR. 2021. ICTV virus taxonomy profile: ovaliviridae. Journal of General Virology 102:001546.

21. Prangishvili D, Liu Y, Krupovic M, Consortium IR. 2021. ICTV virus taxonomy profile: portogloboviridae. Journal of General Virology 102:001605.

22. Prangishvili D, Mochizuki T, Krupovic M, Consortium IR. 2020. ICTV virus taxonomy profile: spiraviridae. Journal of General Virology 101:240–241.

23. Erdmann S, Tschitschko B, Zhong L, Raftery MJ, Cavicchioli R. 2017. A plasmid from an Antarctic haloarchaeon uses specialized membrane vesicles to disseminate and infect plasmid-free cells. Nat Microbiol 2:1446–1455.

24. Al-Shayeb B, Schoelmerich MC, West-Roberts J, Valentin-Alvarado LE, Sachdeva R, Mullen S, Crits-Christoph A, Wilkins MJ, Williams KH, Doudna JA. 2022. Borgs are giant genetic elements with potential to expand metabolic capacity. Nature 610:731–736.

25. Valentin-Alvarado LE, Shi L-D, Appler KE, Crits-Christoph A, De Anda V, Adler BA, Cui ML, Ly L, Leão P, Roberts RJ. 2024. Complete genomes of Asgard archaea reveal diverse integrated and mobile genetic elements. Genome Research 34:1595–1609.

26. Wu F, Speth DR, Philosof A, Cremiere A, Narayanan A, Barco RA, Connon SA, Amend JP, Antoshechkin IA, Orphan VJ. 2022. Unique mobile elements and scalable gene flow at the prokaryote-eukaryote boundary revealed by circularized Asgard archaea genomes. Nat Microbiol 7:200–212.

27. Hou X, He Y, Fang P, Mei SQ, Xu Z, Wu WC, Tian JH, Zhang S, Zeng ZY, Gou QY, Xin GY, Le SJ, Xia YY, Zhou YL, Hui FM, Pan YF, Eden JS, Yang ZH, Han C, Shu YL, Guo D, Li J, Holmes EC, Li ZR, Shi M. 2024. Using artificial intelligence to document the hidden RNA virosphere. Cell 187:6929–6942 e16.

28. Tisza MJ, Pastrana DV, Welch NL, Stewart B, Peretti A, Starrett GJ, Pang YS, Krishnamurthy SR, Pesavento PA, McDermott DH, Murphy PM, Whited JL, Miller B, Brenchley J, Rosshart SP, Rehermann B, Doorbar J, Ta’ala BA, Pletnikova O, Troncoso JC, Resnick SM, Bolduc B, Sullivan MB, Varsani A, Segall AM, Buck CB. 2020. Discovery of several thousand highly diverse circular DNA viruses. Elife 9:1–26.

29. Wu Z, Liu S, Ni J. 2024. Metagenomic characterization of viruses and mobile genetic elements associated with the DPANN archaeal superphylum. Nat Microbiol 9:3362–3375.

30. Kieft K, Anantharaman K. 2022. Virus genomics: what is being overlooked? Curr Opin Virol 53:101200.

31. Diemer GS, Stedman KM. 2012. A novel virus genome discovered in an extreme environment suggests recombination between unrelated groups of RNA and DNA viruses. Biology direct 7:13.

32. Gaia M, Meng L, Pelletier E, Forterre P, Vanni C, Fernandez-Guerra A, Jaillon O, Wincker P, Ogata H, Krupovic M, Delmont TO. 2023. Mirusviruses link herpesviruses to giant viruses. Nature 616:783–789.

33. Luque A, Benler S, Lee DY, Brown C, White S. 2020. The missing tailed phages: prediction of small capsid candidates. Microorganisms 8:1–18.

34. Kirchberger PC, Ochman H. 2023. Microviruses: A World Beyond phi X174. Ann Rev Virol 10:99–118.

35. Kirchberger PC, Martinez ZA, Ochman H. 2022. Organizing the global diversity of microviruses. MBio 13:1–17.

36. Bartlau N, Wichels A, Krohne G, Adriaenssens EM, Heins A, Fuchs BM, Amann R, Moraru C. 2021. Highly diverse flavobacterial phages isolated from North Sea spring blooms. ISME J 16:555–568.

37. Holmfeldt K, Solonenko N, Shah M, Corrier K, Riemann L, VerBerkmoes NC, Sullivan MB. 2013. Twelve previously unknown phage genera are ubiquitous in global oceans. Proc Natl Acad Sci U S A 110:12798–12803.

38. Rinke C, Rubino F, Messer LF, Youssef N, Parks DH, Chuvochina M, Brown M, Jeffries T, Tyson GW, Seymour JR, Hugenholtz P. 2019. A phylogenomic and ecological analysis of the globally abundant Marine Group II archaea (Ca. Poseidoniales ord. nov.). ISME J 13:663–675.

39. Coutinho FH, Rosselli R, Rodríguez-Valera F. 2019. Trends of microdiversity reveal depth-dependent evolutionary strategies of viruses in the Mediterranean. Msystems 4:10.1128/msystems.00554-19.

40. Yilmaz S, Allgaier M, Hugenholtz P. 2010. Multiple displacement amplification compromises quantitative analysis of metagenomes. Nat Methods 7:943–944.

41. Anton BP, Roberts RJ. 2021. Beyond Restriction Modification: Epigenomic Roles of DNA Methylation in Prokaryotes. Annu Rev Microbiol 75:129–149.

42. van Kempen M, Kim SS, Tumescheit C, Mirdita M, Lee J, Gilchrist CLM, Soding J, Steinegger M. 2024. Fast and accurate protein structure search with Foldseek. Nat Biotechnol 42:243–246.

43. Edwards H, Abeln S, Deane CM. 2013. Exploring fold space preferences of new-born and ancient protein superfamilies. PLoS Comput Biol 9:e1003325.

44. Holm L. 2020. Using Dali for protein structure comparison. Methods Mol Biol 2112:29–42.

45. Moi D, Bernard C, Steinegger M, Nevers Y, Langleib M, Dessimoz C. 2023. Structural phylogenetics unravels the evolutionary diversification of communication systems in gram-positive bacteria and their viruses. BioRxiv:2023.09. 19.558401.

46. Vernette C, Lecubin J, Sánchez P, Sunagawa S, Delmont TO, Acinas SG, Pelletier E, Hingamp P, Lescot M. 2022. The Ocean Gene Atlas v2. 0: online exploration of the biogeography and phylogeny of plankton genes. Nucleic Acids Research 50:W516–W526.

47. Sunagawa S, Coelho LP, Chaffron S, Kultima JR, Labadie K, Salazar G, Djahanschiri B, Zeller G, Mende DR, Alberti A. 2015. Structure and function of the global ocean microbiome. Science 348:1261359.

48. Ibarbalz FM, Henry N, Brandão MC, Martini S, Busseni G, Byrne H, Coelho LP, Endo H, Gasol JM, Gregory AC. 2019. Global trends in marine plankton diversity across kingdoms of life. Cell 179:1084–1097. e21.

49. Sunagawa S, Acinas SG, Bork P, Bowler C, Eveillard D, Gorsky G, Guidi L, Iudicone D, Karsenti E. 2020. Tara Oceans: towards global ocean ecosystems biology. Nature Reviews Microbiology 18:428–445.

50. Biller SJ, Ryan MG, Li J, Burger A, Eppley JM, Hackl T, DeLong EF. 2025. Distinct horizontal gene transfer potential of extracellular vesicles versus viral-like particles in marine habitats. Nat Commun 16:2126.

51. Linney MD, Eppley JM, Romano AE, Luo E, DeLong EF, Karl DM. 2022. Microbial sources of exocellular DNA in the ocean. Applied and Environmental Microbiology 88:e02093–21.

52. Brum JR, Ignacio-Espinoza JC, Roux S, Doulcier G, Acinas SG, Alberti A, Chaffron S, Cruaud C, De Vargas C, Gasol JM. 2015. Patterns and ecological drivers of ocean viral communities. Science 348:1261498.

53. Chandler M, De La Cruz F, Dyda F, Hickman AB, Moncalian G, Ton-Hoang B. 2013. Breaking and joining single-stranded DNA: the HUH endonuclease superfamily. Nat Rev Microbiol 11:525–538.

54. Camargo AP, Nayfach S, Chen IA, Palaniappan K, Ratner A, Chu K, Ritter SJ, Reddy TBK, Mukherjee S, Schulz F, Call L, Neches RY, Woyke T, Ivanova NN, Eloe-Fadrosh EA, Kyrpides NC, Roux S. 2022. IMG/VR v4: an expanded database of uncultivated virus genomes within a framework of extensive functional, taxonomic, and ecological metadata. Nucleic Acids Res 50:1–11.

55. Camargo AP, Call L, Roux S, Nayfach S, Huntemann M, Palaniappan K, Ratner A, Chu K, Mukherjeep S, Reddy TBK, Chen IA, Ivanova NN, Eloe-Fadrosh EA, Woyke T, Baltrus DA, Castaneda-Barba S, de la Cruz F, Funnell BE, Hall JPJ, Mukhopadhyay A, Rocha EPC, Stalder T, Top E, Kyrpides NC. 2024. IMG/PR: a database of plasmids from genomes and metagenomes with rich annotations and metadata. Nucleic Acids Res 52:D164–D173.

56. Reiter W-D, Palm P, Yeats S. 1989. Transfer RNA genes frequently serve as integration sites for prokaryotic genetic elements. Nucleic acids research 17:1907–1914.

57. Penades JR, Seed KD, Chen J, Bikard D, Rocha EPC. 2025. Genetics, ecology and evolution of phage satellites. Nat Rev Microbiol doi:10.1038/s41579-025-01156-z.

58. Hou P, Zhou R-Q, Jiang Y-L, Yu R-C, Du K, Gan N, Ke F, Zhang Q-Y, Li Q, Zhou C-Z. 2025. Cryo-EM structure of cyanopodophage A4 reveals a pentameric pre-ejectosome in the double-stabilized capsid. Proceedings of the National Academy of Sciences 122:e2423403122.

59. Terzian P, Olo Ndela E, Galiez C, Lossouarn J, Perez Bucio RE, Mom R, Toussaint A, Petit MA, Enault F. 2021. PHROG: families of prokaryotic virus proteins clustered using remote homology. NAR Genom Bioinform 3:lqab067.

60. Pereira O, Hochart C, Auguet J, Debroas D, Galand P. 2019. Genomic ecology of Marine Group II, the most common marine planktonic Archaea across the surface ocean. MicrobiologyOpen 8, e00852.

61. Lucking D, Alarcon-Schumacher T, Erdmann S. 2023. Distribution and Implications of Haloarchaeal Plasmids Disseminated in Self-Encoded Plasmid Vesicles. Microorganisms 12.

62. Fogarty EC, Schechter MS, Lolans K, Sheahan ML, Veseli I, Moore RM, Kiefl E, Moody T, Rice PA, Yu MK, Mimee M, Chang EB, Ruscheweyh HJ, Sunagawa S, McLellan SL, Willis AD, Comstock LE, Eren AM. 2024. A cryptic plasmid is among the most numerous genetic elements in the human gut. Cell 187:1206–1222 e16.

63. Caulton SG, Lambert C, Tyson J, Radford P, Al-Bayati A, Greenwood S, Banks EJ, Clark C, Till R, Pires E, Sockett RE, Lovering AL. 2024. Bdellovibrio bacteriovorus uses chimeric fibre proteins to recognize and invade a broad range of bacterial hosts. Nat Microbiol 9:214–227.

64. Prabhu A, Zaugg J, Chan CX, McIlroy SJ, Rinke C. 2025. Insights Into Phylogeny, Diversity and Functional Potential of Poseidoniales Viruses. Environmental Microbiology 27:e70017.

65. Philosof A, Yutin N, Flores-Uribe J, Sharon I, Koonin EV, Béjà O. 2017. Novel abundant oceanic viruses of uncultured marine group II Euryarchaeota. Current Biology 27:1362–1368.

66. Fujisawa H, Morita M. 1997. Phage DNA packaging. Genes to Cells 2:537–545.

67. Lucía-Sanz A, Manrubia S. 2017. Multipartite viruses: adaptive trick or evolutionary treat? NPJ Systems Biology and Applications 3:34.

68. Alarcón-Schumacher T, Lücking D, Erdmann S. 2023. Revisiting evolutionary trajectories and the organization of the Pleolipoviridae family. PLoS Genetics 19:e1010998.

69. Mishra A, Krishnan B, Srivastava SS, Sharma Y. 2014. Microbial betagamma-crystallins. Prog Biophys Mol Biol 115:42–51.

70. Krupovic M, Koonin EV. 2017. Multiple origins of viral capsid proteins from cellular ancestors. Proc Natl Acad Sci U S A 114:E2401–E2410.

71. Chen S, Zhou Y, Chen Y, Gu J. 2018. fastp: an ultra-fast all-in-one FASTQ preprocessor. Bioinformatics 34:i884–i890.

72. Li D, Liu C-M, Luo R, Sadakane K, Lam T-W. 2015. MEGAHIT: an ultra-fast single-node solution for large and complex metagenomics assembly via succinct de Bruijn graph. Bioinformatics 31:1674–1676.

73. Bouras G, Nepal R, Houtak G, Psaltis AJ, Wormald PJ, Vreugde S. 2023. Pharokka: a fast scalable bacteriophage annotation tool. Bioinformatics 39.

74. Altschul SF, Gish W, Miller W, Myers EW, Lipman DJ. 1990. Basic local alignment search tool. Journal of molecular biology 215:403–410.

75. Abramson J, Adler J, Dunger J, Evans R, Green T, Pritzel A, Ronneberger O, Willmore L, Ballard AJ, Bambrick J. 2024. Accurate structure prediction of biomolecular interactions with AlphaFold 3. Nature 630:493–500.

76. Meng EC, Goddard TD, Pettersen EF, Couch GS, Pearson ZJ, Morris JH, Ferrin TE. 2023. UCSF ChimeraX: Tools for structure building and analysis. Protein Science 32:e4792.

77. Sievers F, Wilm A, Dineen D, Gibson TJ, Karplus K, Li W, Lopez R, McWilliam H, Remmert M, Söding J. 2011. Fast, scalable generation of high-quality protein multiple sequence alignments using Clustal Omega. Mol Syst Biol 7:1–7.

78. Eddy SR. 2011. Accelerated profile HMM searches. PLoS Comput Biol 7:e1002195.

79. Togkousidis A, Kozlov OM, Haag J, Hohler D, Stamatakis A. 2023. Adaptive RAxML-NG: Accelerating Phylogenetic Inference under Maximum Likelihood using Dataset Difficulty. Mol Biol Evol 40.

